# Hyperexcitability and translational phenotypes in a preclinical model of *SYNGAP1* mutations

**DOI:** 10.1101/2023.07.24.550093

**Authors:** Timothy A Fenton, Olivia Y Haouchine, Elizabeth L Hallam, Emily M Smith, Kiya C. Jackson, Darlene Rahbarian, Cesar Canales, Anna Adhikari, Alexander S. Nord, Roy Ben-Shalom, Jill L Silverman

## Abstract

*SYNGAP1* is a critical gene for neuronal development, synaptic structure, and function. Although rare, the disruption of *SYNGAP1* directly causes a genetically identifiable neurodevelopmental disorder (NDD) called SYNGAP1-related intellectual disability. Without functional SynGAP1 protein, patients present with intellectual disability, motor impairments, and epilepsy. Previous work using mouse models with a variety of germline and conditional mutations has helped delineate SynGAP1’s critical roles in neuronal structure and function, as well as key biochemical signaling pathways essential to synapse integrity.

Homozygous loss of *SYNGAP1* is embryonically lethal. Heterozygous mutations of *SynGAP1* result in a broad range of phenotypes including increased locomotor activity, impaired working spatial memory, impaired cued fear memory, and increased stereotypic behavior. Our *in vivo* functional data, using the original germline mutation mouse line from the Huganir laboratory, corroborated robust hyperactivity and learning and memory deficits. Here, we describe impairments in the translational biomarker domain of sleep, characterized using neurophysiological data collected with wireless telemetric electroencephalography (EEG). We discovered *Syngap1*^+/-^ mice exhibited elevated spike trains in both number and duration, in addition to elevated power, most notably in the delta power band. Primary neurons from *Syngap1*^+/-^ mice displayed increased network firing activity, greater spikes per burst, and shorter inter-burst intervals between peaks using high density micro-electrode arrays (HD-MEA). This work is translational, innovative, and highly significant as it outlines functional impairments in *Syngap1* mutant mice. Simultaneously, the work utilized untethered, wireless neurophysiology that can discover potential biomarkers of Syngap1R-ID, for clinical trials, as it has done with other NDDs. Our work is substantial forward progress toward translational work for SynGAP1R-ID as it bridges *in-vitro* electrophysiological neuronal activity and function with *in vivo* neurophysiological brain activity and function. These data elucidate multiple quantitative, translational biomarkers *in vivo* and *in vitro* for the development of treatments for SYNGAP1-related intellectual disability.

## Introduction

Many severe neurodevelopmental disorders (NDDs) include underlying excitatory/inhibitory imbalances and seizures. These underlying imbalances are thought to worsen behavioral symptoms of NDDs, such as autism spectrum disorders (ASD) and intellectual disabilities (ID) and underlie cognitive decline and impaired cognitive development. The *SYNGAP1* (synaptic Ras GTPase activating protein) gene encodes the protein (SynGAP1), which is selectively expressed in the brain, and highly enriched at excitatory synapses,^1, 2^ and is critical for neuronal development and synaptic plasticity.^3^

Detailed research on SynGAP since its description in 1998 has produced high quality data on its own protein structure, role in neuronal structure, biochemical and physiological function and its unique neuronal localization.^4, 5^ Reduction or loss of SynGAP1 leads to Ras activation and excessive AMPA receptor incorporation into the cell membrane,^6^ components critical for long[term potentiation, dendritic spine formation, neuronal development and structural integrity, neuronal signaling,^3^ synaptic strength or dysregulation,^4^ and the long term potentiation processes that underlie cognition and excitability.^5^

SYNGAP1-related intellectual disability (SYNGAP1R-ID) is an NDD characterized by global developmental delay, ASD, ID, and epilepsy. In most individuals, the ID is moderate to severe, and the epilepsy is either generalized or has myoclonic absence events. Loss of this crucial function at the synapse results in dysfunctional and aberrant neuronal signaling. SYNGAP1R-ID can therefore be considered a ‘synaptopathy’, which refers to disorders caused by synaptic dysfunction that leads to aberrant neuronal communication.^7^ In addition to the dysregulation of synaptic signaling, loss of SynGAP1 also results in aberrant Ras GTPase cellular signaling, making SynGAP1R-ID, a Rasopathy. The ‘Rasopathies’ refer to a group of brain disorders in which the RAS/MAPK signaling pathway is disrupted. This dual dysregulation of inter- and intracellular communication results in debilitating and severe consequences clinically. De novo *SYNGAP1* mutations have been found in patients with developmental delays and ID (96%), epilepsy (98%), and/or ASD (50%).^8, 9^

As SynGAP1 is a negative regulator of excitatory neurotransmission, overexpression of SynGAP1 results in a dramatic loss of synaptic efficacy. Conversely, enhanced synaptic transmission occurs when SynGAP1 is disrupted by RNA interference,^4^ highlighting the fact that SynGAP1 is critical for multiple processes, given its essential role at the synapse and in cellular signaling. As described by Rumbaugh et al., SynGAP1 is modifiable with a variety of modern technologies, so restoration of SynGAP1 is not a hopeless endeavor. Targeted molecular strategies and therapeutics, including adeno associated viral vectors,^9^ CRISPRa activation,^10^ activating RNAs,^11^ and antisense oligonucleotides (ASOs)^12^ are in development. In addition to precision therapeutics, SYNGAP1R-ID, being adjacent to Rasopathies and Synaptopathies, widens opportunities to repurpose traditional pharmacologic compounds.

Given the outstanding need to develop effective therapies for SYNGAP1R-ID, our laboratory has been focused on biomarkers and functional outcome measures that are rigorous, robust, reliable, and objective. Critically, herein, we provide a seminal report of behavioral impairments resulting from the loss of SynGAP1, translational biomarkers of sleep and power spectral density bands using EEG. These data confirm a unique electrophysiological signature in live *Syngap1* mutant mice as well as in cultured primary neurons, bridging electrophysiological single neuronal network firing patterns *in vitro* to neurophysiological and behavioral phenotypes *in vivo*. Our HD-MEA work here is groundbreaking and paves the way for functional outcomes that bridge mouse neurons to neural stem cells, reprogrammed from human iPSCs.

## Methods

### Animals

All animals were housed in Techniplast cages (Techniplast, West Chester, PA, USA) in a temperature (68-72°F) and humidity (∼25%) controlled vivarium maintained on a 12:12 light-dark cycle. All procedures were approved by the Institutional Animal Care and Use Committee at the University of California Davis and were conducted in accordance with the National Institutes of Health Guide for the Care and Use of Laboratory Animals. B6;129-*Syngap1*^tm1Rlh^/J mice were obtained from The Jackson Laboratory (JAX #008890, Bar Harbor, ME, USA) and fed a standard diet of Teklad global 18% protein rodent diets 2918 (Envigo, Hayward, CA, USA). Rodent chow and tap water were available *ad libitum*. In addition to standard bedding, a Nestlet square, shredded brown paper, and a cardboard tube (Jonesville Corporation, Jonesville, MI) were provided in each cage. Heterozygous deletion male mice were bred with 129S1-C57BL/6J F1 hybrid females to generate mutant (*Syngap1^+/-^*) and wildtype (WT, *Syngap1^+/+^*) littermates. The 129S1-C57BL/6J F1 hybrid mice are the offspring of a cross between C57BL/6J females and 129S1/SvImJ males (JAX #101043). To identify mice, pups were labeled by paw tattoo on postnatal days (PND) 2–4 using non-toxic animal tattoo ink (Ketchum Manufacturing Inc., Brockville, ON, Canada). Tails of pups were clipped (1–2 mm) for genotyping following the UC Davis IACUC policy regarding tissue collection. Genotyping was performed with REDExtract-N-Amp (Sigma Aldrich, St. Louis, MO, USA) using primers JAX oIMR9462 ATGCTCCAGACTGCCTTGGGAAAAG, oIMR9463 ACCTCAAATCCACACTCCTCTCCAG, and oIMR9464 AGGGAACATAAGTCTTGGCTCTGTC.

To reduce carry-over effects from repeated behavioral testing, at least 24 hours were allowed to pass between the completion of one task and the start of another. Assays were performed in the order of least to most stressful and between the hours of 8:00AM PST and 7:00PM PST during the light phase. All behavior testing was conducted by an experimenter blinded to genotype and included both sexes. Mice were allowed to habituate in their home cages in a dimly lit room adjacent to the testing room for 1 hour prior to the start of testing to limit any effect of transporting between the vivarium and testing suite. Between subjects, testing apparatus surfaces were cleaned using 70% ethanol and allowed to dry. One cohort of animals was tested comprised of 13 litters beginning at 8 weeks of age (postnatal day (PND) 55). The order of testing was (1) elevated plus-maze, (2) light ↔ dark transitions test, (3) open field, (4) spontaneous alternation, (5) novel object recognition.

### Immunohistochemistry and Imaging

Following anesthesia with isoflurane, Postnatal day 60 (PND60) mice were transcardially perfused with 4% paraformaldehyde (PFA) in 1xPBS and brains post-fixed overnight in the same solution at 4°C. Brains were then transferred to 30% sucrose/PBS solution to equilibrate until they sank to the bottom of a conical tube. Once equilibrated, OCT-embedded (Tissue-Tek) brains were cryo-sectioned coronally (30µm) and collected for free-floating immunostaining with agitation. Sections underwent antigen retrieval using 1x Citrate buffer pH6.0 antigen retriever solution (C9999, Millipore-Sigma, Burlington, MA), at 60°C for 1 hour. Subsequently, sections were permeabilized in PBS containing 0.5% Triton X-100 for 20 minutes and incubated in blocking solution for 1 hour at room temperature in 5% milk/PBST (1x PBS with 0.1% Triton X-100). The SynGAP1 immunolabeling was performed using a commercially available Anti-SynGAP1 antibody (1:250; PA1-046; ThermoFisher Scientific), incubating overnight at 4°C with orbital agitation. A no-primary antibody control was used to evaluate specificity. After primary antibody incubation, free-floating sections were washed five times with PBST (20 minutes each). Species-specific fluorophores-conjugated IgG (1:500; Thomas Scientific) was used as secondary antibodies (45 minutes, RT). 40,6-Diamidion-2-phenylindole (DAPI; 1:1000) was used for nuclear staining (25 min, RT). Imaging was carried out on a Keyence all-in-one fluorescence (BZ-X810). FIJI (National Institutes of Health) was used for image processing with settings consistently applied across samples.

### Protein extraction and western blot analysis

Brains were extracted from PND42 mice. The cortex was collected then snap-frozen on dry ice for storage at -80[. The cortical tissue was suspended in 600μl lysis buffer (50mM Tris-HCl, 140mM NaCl, 10% Glycerol, 0.5% IGEPAL, 0.25% Triton X-100, protease inhibitor cocktail (Roche, 4693124001)) and manually homogenized. After a 30-minute incubation on ice, samples were lysed with a probe sonicator (Qsonica CL-18, 20% amplitude, 10 cycles of 5sec on/off intervals) and placed back on ice to incubate for another 30 minutes. Cell debris was collected by centrifugation (14000 g, 4[, 10min) and the supernatant was moved to a new tube. Protein concentration was quantified using a BCA protein assay kit (Pierce, 23225) and samples were stored at -80[ until use.

Protein lysates were diluted in 30μl 6X Laemmli SDS buffer (375mm Tris-HCl, 9% SDS, 50% glycerol, 0.03% Bromophenol blue) and 5% β-mercaptoethanol, boiled at 70[ for 10 minutes, and separated on 4-20% polyacrylamide tris-glycine protein gel (BioRad, 4561094). The separated proteins were transferred onto a PVDF membrane (Millipore Sigma, 0.45μm, IPFL00010) by wet transfer (overnight, constant 13mA at max voltage, 4[). Membranes were blocked with Intercept PBS blocking buffer (Li-Cor, LIC-927-90001) at room temperature for one hour. Primary antibodies were diluted in 7.5ml Intercept PBS blocking buffer with 0.1% Tween (SynGAP1; 1:1000dil, Invitrogen, PA1-046, Gapdh; 1:15000, Sigma-Aldrich G8795). Membranes were incubated with the primary antibody solution overnight at 4[, then washed four times for 5 minutes with PBS with 0.1% Tween (PBST). Fluorescently tagged secondary antibodies (Li-Cor, 926-32212 and 926-68023) were diluted in 10ml Intercept PBS blocking buffer with 0.1% Tween. After the initial washes, blots were incubated with the secondary antibody solution for one hour at room temperature. Blots were washed an additional four times for 5 minutes with PBST and two times with PBS. Bands were visualized using the Odyssey DLx imaging system (Li-Cor).

### Elevated-plus maze

The elevated-plus maze (EPM) is a well-established task for assessing anxiety-like conflict behavior in rodents by allowing mice to choose between entering the two open arms of the maze (natural exploratory drive) or entering and remaining in the safety of the two closed arms. All four arms are elevated 1 m from the floor, with the drop-off detectable only in the open arms. The EPM was performed according to previously described procedures^13–16^ using a mouse EPM (model ENV-560A) purchased from Med Associates (St. Albans, VT). The EPM contained two open arms (35.5 cm × 6 cm) and two closed arms (35.5 cm × 6 cm) radiating from a central area (6 cm × 6 cm). A 0.5 cm high lip surrounded the edges of the open arms, whereas the closed arms were surrounded by 20 cm high walls. The EPM was cleaned with 70% ethanol before the beginning of the first test session and after each subject mouse was tested, with sufficient time for the ethanol to dry and for the odor to dissipate before the start of the next test session. The room was illuminated at ∼ 40 lx. To begin the test, the mouse was placed in the central area facing the open arm. The mouse was allowed to freely explore for 5 min during which time the activity was recorded by a computer counting beam breaks between arms.

### Light ↔ dark transitions

The light ↔ dark transitions test assesses anxiety-like conflict behavior in mice by evaluating the tendency of mice to avoid brightly lit areas versus their strong tendency to explore a novel environment. The light ↔ dark transitions test was performed in accordance with previously described procedures.^13–16^ The test began by placing the mouse in the light side (∼ 320 lx; 28 cm × 27.5 cm × 27 cm) of an automated 2-chambered apparatus, in which the enclosed/dark side (∼ 5 lx; 28 cm × 27.5 cm × 19 cm) was reached by traversing the small opening of the partition between the two chambers. The mouse was allowed to explore freely for 10 min. Time in the dark side chamber and total number of transitions between the light and dark side chambers were automatically recorded during the 10-minute session using Labview 8.5.1 software (National Instruments, Austin, TX).

### Open Field

General exploratory locomotion in a novel open field environment was assayed in an arena sized 40 cm × 40 cm × 30.5 cm, as previously described.^13–15, 17–22^ Open field activity was considered an essential control for effects on physical activity, for example, as sedation or hyperactivity could confound the interpretation of interaction time with an arena or objects. The testing room was illuminated at ∼ 40 lx.

### Spontaneous Alternation

Spontaneous alternation in a Y-maze was assayed using methods modified from previous studies.^13, 14, 17^ The Y-maze assesses working learning and memory and has been validated by experiments in which scopolamine (0.56 and 1 mg/kg; 30 min pre, i.p.) produced significant reductions in spontaneous alternation behavior relative to vehicle-treated C57Bl/6J mice (communication with Dr. Sukoff-Rizzo and open access https://www.jax.org/MNBF). Subjects explored a Y-shaped maze constructed of matte white acrylic (P95 White, Tap Plastics, Sacramento, CA, USA) for 8 minutes and were recorded from an overhead camera with the behavioral tracking software Ethovision XT. Mice were placed at the center of the initial arm facing the center of the maze. Percentage of spontaneous alternations is calculated as the number of triads (entry into three different arms without returning to a previously entered arm) relative to the number of alternation opportunities. All scoring was conducted by an observer blinded to genotype.

### Novel Object Recognition

The novel object recognition (NOR) test was conducted in opaque matte white (P95 White, Tap Plastics, Sacramento, CA) open field arenas (41 cm × 41 cm × 30cm), using methods similar to those previously described.^14, 16^ The experiment consisted of 4 sessions: a 30-min exposure to the open field arena the day before the test, a 10-min re-habituation on test day, a 10-min familiarization session and a 5-min recognition test. On day 1, each subject was habituated to a clean, empty open field arena for 30-min. 24-hr later, each subject was returned to the open field arena for another 10-min for the habituation phase. The mouse was then removed from the open field and placed in a clean temporary holding cage for approximately 2-min. Two identical objects were placed in the arena. Each subject was returned to the open field in which it had been habituated and allowed to freely explore for 10-min. After the familiarization session, subjects were returned to their holding cages, which were transferred from the testing room to a nearby holding area. The open field was cleaned with 70% ethanol and let dry. One clean familiar object and one clean novel object were placed in the arena, where the two identical objects had been located during the familiarization phase. 60-min after the end of the familiarization session, each subject was returned to its open field for a 5-min recognition test, during which time it was allowed to freely explore the familiar object and the novel object. The familiarization session and the recognition test were recorded with Ethovision XT video tracking software and manually scored by an experimenter blinded to genotype (Version 9.0, Noldus Information Technologies, Leesburg, VA). Object investigation was defined as time spent sniffing the object when the nose was oriented toward the object and the nose–object distance was 2-cm or less. Recognition memory was defined as spending significantly more time sniffing the novel object compared to the familiar object via a within group paired t-test. Total time spent sniffing both objects was used as a measure of general exploration. Time spent sniffing two identical objects during the familiarization phase confirmed the lack of an innate side bias. Objects used were plastic toys: a small soft plastic orange safety cone and a hard plastic magnetic cone with ribbed sides, as previously described.^13, 14, 17, 23, 24^

### Electroencephalography (EEG)

A cohort of 19 animals were surgically implanted with wireless EEG telemetric devices (HD-X02, Data Sciences International, St. Paul, MN, USA), as previously described.^13, 18, 19, 25^ The implantation procedure was performed in accordance with the UC Davis IACUC Guidelines for Rodent Survival Surgery. All mice aged 2-4 months old and weighing over 20g, were anesthetized with vaporized liquid isoflurane (Piramal Critical Care, Inc., Bethlehem, PA, USA).

With a micro drill (Stoelting), two 1-mm-diameter burr holes were manually drilled (1.0 mm anterior and 1.0 mm lateral; − 3.0 mm posterior and 1.0 mm lateral relative to bregma) allowing for the placement of two steel surgical screws (00-96 X 1/16 IROX screw, DSI, MN, USA). A subcutaneous pocket lateral to the spine was then made using a Crile Hemostat, minimizing excess tissue damage, and avoiding potential discomfort to the animal once implanted. Attached to the implant were two pairs of reference and sensing leads made of a nickel cobalt-based alloy insulated with medical-grade silicone, used to collect EEG and EMG biopotential data.

One set of leads, used to measure EEG activity across the frontal cortical area, were individually attached to a surgical screw by removing the silicone insulation from the terminal end of the lead and tying the exposed wire around the base of the screw. The remaining set of leads were used to measure EMG activity.

Animals were housed in a temperature controlled ventilated cabinet (Aria Bio-C36 EVO Ventilated Cabinet, Techniplast, Maggio, Italy) to support the maintenance of core body temperature by limiting activity dependent thermoregulation to improve postoperative recovery. Analgesic (carprofen) was administered the day after surgery, and as necessary following a thorough health evaluation performed twice a day during the 7-10 day postoperative period.

Untethered EEG activity was recorded in the home cage of individually housed, freely moving mice each assigned to a PhysioTelTM RPC-1 receiver plate (DSI, MN, USA). EEG and EMG data were collected at a sampling rate of 500 Hz with a 0.1 Hz high-pass and 100 Hz low-pass bandpass filter. The signal was transmitted to a control box which facilitates the transformation of signal between each implant and receiver pairing to the acquisition computer running Ponemah software (DSI, MN, USA). Activity, temperature, and signal strength were collected at a sampling rate of 200 Hz.

### Cell Culture

Primary cortical neuron-glia cultures were prepared using brain tissue from PND0-1 *Syngap1^+/-^* pups as previously described.^26^ Briefly, cortices were minced with a razor blade prior to incubation in Hibernate A containing papain and DNase. The tissue was then triturated with glass pipettes to dissociate cells from the extracellular matrix (Bellco Glass, Vineland, NJ). Dissociated cells were counted and plated on microelectrode array chips precoated with polyethyleneimine (PEI) (Sigma-Aldrich) and laminin (Sigma-Aldrich).

### Micro-Electrode Array Electrophysiology (MEA)

MaxWell Biosystems high density microelectrode arrays (HD-MEAs) were used to assess electrophysiological activity of primary neuronal cultures. HD-MEA chips were incubated with a 1% Tergazyme solution (Alconox, White Plains, NY) for 2 hours at room temperature then washed with DI H_2_O before being transferred to a beaker of 70% ethanol. The beaker containing the HD-MEA chips was transferred to the biosafety cabinet and allowed to incubate for 30 minutes at room temperature to sterilize the chips. HD-MEA chips were then washed with sterile water three times before adding 1 ml of complete cell culture media consisting of Neurobasal supplemented with B27 (Gibco, ThermoFisher Scientific) and 10% horse serum (Gibco, ThermoFisher Scientific). The HD-MEA chips were allowed to incubate for two days in a humidified cell culture incubator at 37C with 5% CO_2_. On the day of cell plating, the cell culture media was removed, and chips were washed three times with sterile water before being coated with 50µl of polyethyleneimine (PEI, Sigma). Chips were placed back in the incubator for one hour, then the coating was aspirated, and the chips were washed three times with sterile water and allowed to dry in the biosafety cabinet for one hour. Laminin was then applied to the chips and placed back in the incubator until ready to plate cells (∼1 hr). The laminin coating was removed and 50µl of primary neuronal cells isolated as previously described was immediately pipetted onto the HD-MEAs resulting in ∼50,000 cells per chip. After incubating for 1 hour in the cell culture incubator, an additional 600µl of cell culture medium was added to each chip. The next day, 50% of the media in each chip was removed and replaced with fresh cell culture media. Media changes happened every three days by removing 50% of the media and replacing it with an equal volume of cell culture media.

HD-MEA recordings were performed between day *in vitro* (DIV) 17 and DIV35, based on previous published literature.^27–29^ The MaxWell Biosystems recording unit was sterilized with 70% ethanol, placed in the biosafety cabinet, and allowed to dry for 30 minutes. The recording unit was then transferred to an incubator at 37°C with 5% CO_2_ for at least two hours prior to recording to allow for temperature equilibration. Performing MEA recordings inside the incubator ensured consistent temperature and pH values of the cell culture media while performing various scans. To assess neuronal electrical activity across the entire electrode array, the “Activity Scan Assay” module in the MaxLab Live software (MaxWell Biosystems AG, Zurich, Switzerland) was used. The “Full Scan” electrode configuration was used to measure neuronal activity from all 26,400 electrodes for 30 seconds each. After the Activity Scan was complete, the “Network Assay” module was used to assess network activity or axonal features. Network electrical activity was recorded by selecting a subset of 1024 electrodes with the highest firing rate from the corresponding chip’s Activity Scan and measured simultaneously for 300 seconds. Custom code was written to analyze the data using MatLab R2021.

### Experimental Design and Statistical Analysis

Data were analyzed with GraphPad Prism. Statistical testing was performed using established assay-specific methods, including Student’s t-test for single parameter comparisons between genotypes, and one-way or two-way repeated-measures ANOVA for comparisons across time points. All significance levels were set at p < 0.05 and all t-tests were two-tailed. Group sizes were chosen based on experience and power analyses.^15, 22, 30^ Significant ANOVAs were followed by Bonferroni-Dunn or Holm-Sidak posthoc testing. Behavioral analysis passed distribution normality tests, was collected using continuous variables and thus was analyzed via parametric analysis in all assays. For all behavioral analyses, variances were similar between groups and data points within two standard deviations of the mean were included in analysis. Finally, sex differences were not observed in any behavioral assay previously and thus sexes were combined.

## Data Availability

The data that support the findings of this study are available from the corresponding author, upon reasonable request.

## Results

### Internal Validation of reduced Syngap1 protein in *Syngap1^+/-^* mice

*SynGAP1*^+/-^ gene mutations reduce the expression of the wild-type SynGAP protein. *Syngap1^+/-^* mice had a significant decrease in the protein expression of SynGAP1 compared to *Syngap1^+/+^* mice. Western blot analysis was performed on brain lysates to measure the level of SynGAP1 protein expression. *Syngap1^+/-^* mice had significantly less SynGAP1 expression when compared to *Syngap1^+/+^* mice at age PND42 (**Fig. 1A**). The heterozygous mutation reduced SynGAP1 protein production to 41% of wildtype expression (**Fig. 1B**; *t*(3) = 7.3937, *P* = 0.0051). A representative mouse brain coronal section from PND60 *Syngap1^+/+^* animal shows broad SynGAP1 protein expression through immunostaining (red). DAPI staining (nuclei, blue) helps identify brain structures. Boxes indicate magnified areas. Scale bars = 100 µm. Immunoblot analysis of mouse brain homogenates at P5 with the anti-GAP SynGAP antibody showed that expression of the 130kDa SynGAP protein is abolished in the mutant mice (**Fig. 1A**). However, overexposure of the immunoblot showed that a very low level of a smear of smaller proteins (120 kDa) could be detectable using the anti-GAP antibody. Broad SynGAP1 protein expression was detectable in coronal brain section from PND60 *Syngap1^+/+^* mice across different brain regions. SynGAP1 was detected in the cytosol in the primary somatosensory area of the cortex and in the CA2 region of the hippocampus when representative high-power fields of observation were evaluated (**Fig. 1C**).

**Figure 1.**
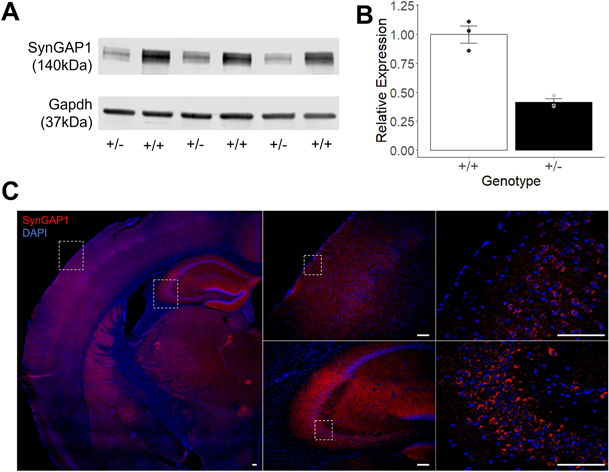
*Syngap1^+/-^*mice had a significant decrease in the protein expression of SynGAP1 compared to *Syngap1^+/+^* mice. **(A)** SynGAP1 and Gapdh protein expression in *Syngap1^+/+^* (WT) and *Syngap1^+/-^* (HT) mice at PND42. Western Blot analysis of SynGAP1 (140kDa) shows a decrease in expression of SynGAP1 in the *Syngap1^+/-^*mice. **(B)** Quantification of SynGAP1 protein expression normalized using Gapdh constitutive expression. SynGAP1 protein expression was significantly decreased to 41% in *Syngap^+/-^*brains as compared to *Syngap1^+/+^* littermates. **(C)** Representative mouse brain coronal section from PND60 *Syngap1^+/+^* animal shows broad SynGAP1 protein expression through immunostaining (red). DAPI staining (nuclei, blue) helps identify brain structures. Boxes indicate magnified areas. Scale bars = 100 µm. Data was analyzed using a Student’s t-test and is expressed as mean ± S.E.M. \**P* = 0.0051. (+/+ *N* = 3, *Syngap^+/-^ N* = 3).

### Increased locomotion in the open field assay

*Syngap1^+/-^* mice displayed hyperactivity in the open field assay. Significantly more locomotion was observed in *Syngap1^+/-^*mice when compared to *Syngap1^+/+^* mice at every time bin of a 30-minute session when horizontal activity was measured and analyzed with a two-way ANOVA and subsequent comparison via Sidak’s post hoc test between genotypes (**Fig. 2A**; *F*(1, 46) = 87.18, *P* < 0.0001 (main effect); 1-5 min, *P* < 0.0001; 6-10 min, *P* < 0.0001; 11-15 min, *P* < 0.0001; 16-20 min, *P* < 0.0001; 21-25 min, *P* < 0.0001; 26-30 min, *P* < 0.0001). Similar elevated levels of activity were observed at every time point when total activity was assessed when compared with a two-way ANOVA between genotypes (**Fig. 2B**; *F*(1, 46) = 88.90, *P* < 0.0001 (main effect); 1-5 min, *P* < 0.0001; 6-10 min, *P* < 0.0001; 11-15 min, *P* < 0.0001; 16-20 min, *P* < 0.0001; 21-25 min, *P* < 0.0001; 26-30 min, *P* < 0.0001). Increased center time was significantly increased in *Syngap1^+/-^* mice when compared to *Syngap1^+/+^* mice at five of the six five-minute time bins when compared with Sidak’s post hoc test (**Fig. 2C**; *F*(5, 230) = 8.573; 6-10 min, *P* = 0.0001; 11-15 min, *P* < 0.0001; 16-20 min, *P* = 0.0003; 21-25 min, *P* = 0.0002; 26-30 min, *P* < 0.0001).

**Figure 2.**
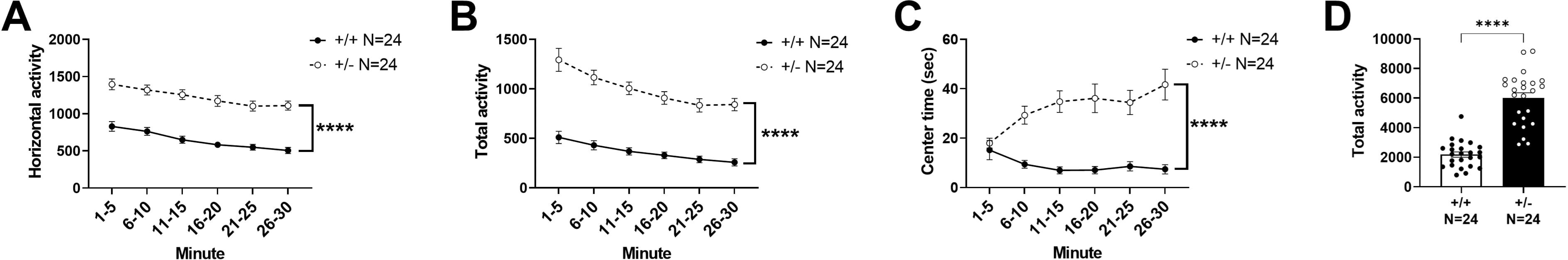
*Syngap1^+/-^*mice showed elevated motor activity in the open field assay. **(A)** Horizontal activity, **(B)** total activity, and **(C)** center time were all significantly increased when compared to *Syngap1^+/+^* mice. Horizontal activity, total activity, and center time were analyzed using a two-way ANOVA. **(D)** Total activity over 30 minutes was compared with a Student’s t-test. Data are expressed as mean ± S.E.M. * = *P* < 0.05, **** = *P* < 0.0001.

Taken together with the increased time spent in the open arms and transitions in the elevated plus maze (**Fig. S1**), the elevated center time indicates a reduced anxiety-like phenotype as well as a hyperactive locomotor phenotype. When summed over the 30-minute session, the total activity of the *Syngap1^+/-^* mice was significantly higher than the *Syngap1^+/+^* mice following a student’s unpaired t-test (**Fig. 2D**; *t*(46) = 9.429, *P* < 0.0001).

### Cognitive assessment in the Novel Object Recognition and Y-maze Tasks

Cognitive abilities were tested in both novel object recognition task (long-term memory) and the Y-maze (working learning and memory). Interestingly, even with a hyperactive phenotype seen in the elevated plus maze, light-dark task, and the open field task, we observed a reduced preference for the novel object in the *Syngap1*^+/-^ mice when compared with a Student’s t-test between familiar and novel objects indicating a cognitive deficit (**Fig. 3A**; *t*(43) = 1.572, *P* = 0.1234). As expected, the *Syngap1^+/+^* mice spent significantly more time investigating the novel object compared to the familiar object when analyzed with a Student’s t-test (**Fig. 3A**; *t*(43) = 2.512, *P* = 0.0158). In the Y-maze, *Syngap1^+/-^* mice did not significantly differ from WT mice in the percentage of triads (**Fig. 3B**; *t*(46) = 0.2335, *P* = 0.8164). However, the *Syngap1^+/-^* mice made significantly more transitions between arms in the Y-maze when compared to *Syngap1^+/+^*mice providing further evidence of their hyperactivity phenotype (**Fig. 3C**; *t*(45) = 8.154, *P* < 0.0001).

**Figure 3.**
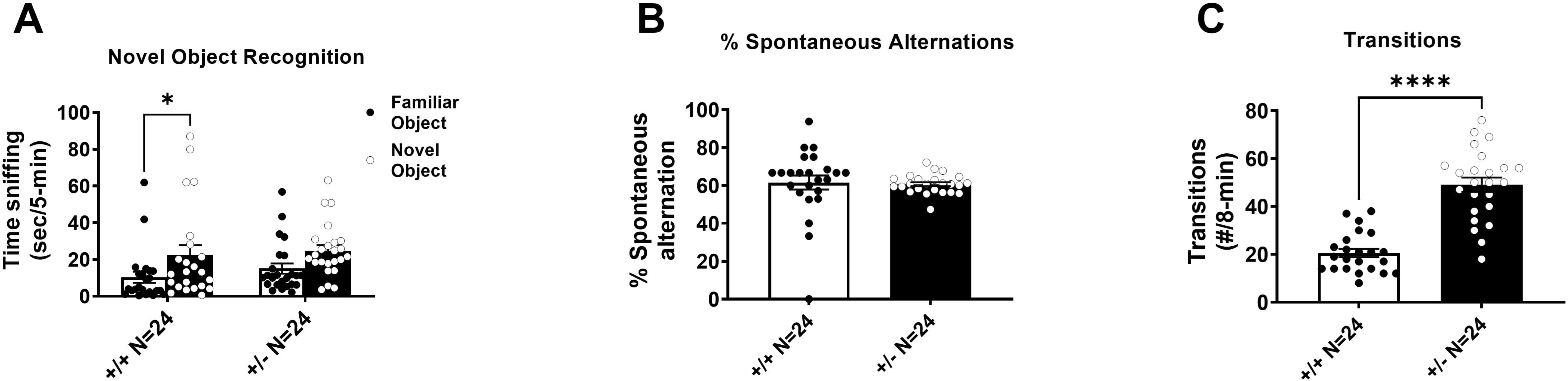
*Syngap1^+/-^*mice showed impaired cognition in the novel object recognition task and hyperactivity in the spontaneous alternation task. **(A)** *Syngap1^+/-^* mice showed no preference for the novel object when compared to wildtype mice. **(B)** In the spontaneous alternation task, there was no difference in the percentage of alternations between wildtype and *Syngap1^+/-^* animals. **(C)** *Syngap1^+/-^*mice made significantly more total transitions between arms in the spontaneous alternation task when compared to wildtype animals. Groups compared with a Student’s t-test. Data are expressed as mean ± S.E.M. * = *P* < 0.05, **** = *P* < 0.0001.

### Altered *in vivo* electroencephalography in *Syngap1^+/-^* mice

A wireless telemeter system was used to measure electroencephalographic activity. *Syngap1^+/-^* mice displayed an elevated absolute power spectral density (PSD) compared to *Syngap1^+/+^*mice when measured for 72 hours and compared with a two-way ANOVA between genotype and frequency (**Fig. 4A**; *F*(1, 13104) = 134.5, *P* < 0.0001). When observing differences in different power bands, *Syngap1^+/-^* mice displayed elevated Delta power (.5-4 HZ) and Theta power (5-9 HZ) when compared with a two-way ANOVA between genotypes and subsequent post hoc analysis (**Fig. 4B**; *F*(1, 16) = 8.598, *P* = 0.0098; *P* < 0.0001 (Delta), *P* = 0.0287 (Theta)). Delta power and theta power bands were elevated in *Syngap1^+/-^*mice in representative total power distributions (**Fig. 4C**). We observed a significantly increased spike train count in *Syngap1^+/-^* mice compared to *Syngap1^+/+^*mice (**Fig. 4D**; *t*(13) = 4.396, *P* = 0.0007). The duration of spike trains was also significantly elevated in *Syngap1^+/-^* mice compared to *Syngap1^+/+^*mice (**Fig. 4E**; *t*(13) = 3.215, *P* = 0.0068). *Syngap1^+/-^* mice displayed higher signal than *Syngap1^+/+^* mice when viewing raw EEG output (**Fig. 4F**).

**Figure 4.**
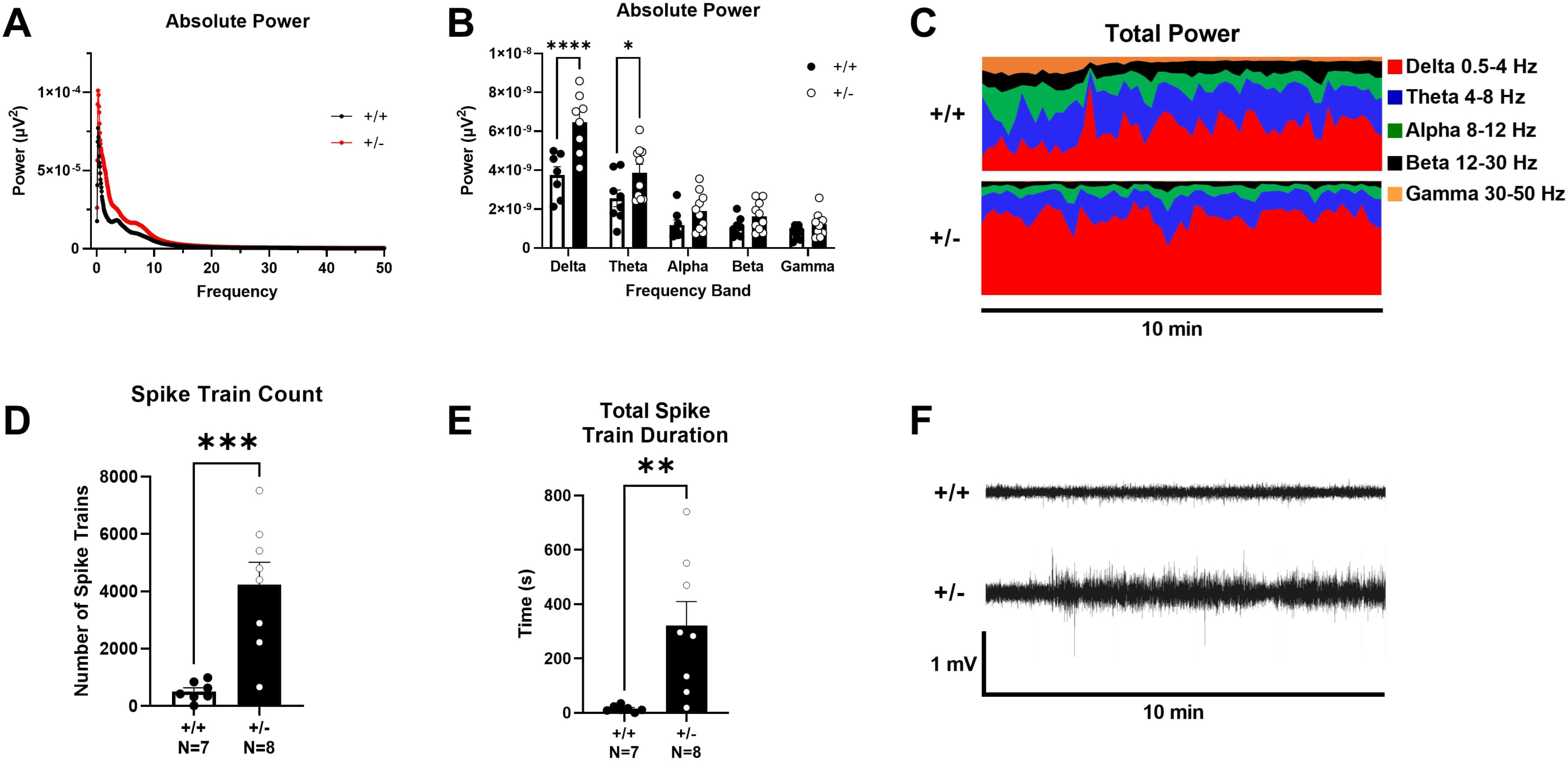
*Syngap1^+/-^*mice displayed elevated delta and theta power. Surface EEG was collected using wireless telemetry and power spectral densities were compared at delta (.5-4 Hz), theta (4-8 Hz), alpha (8-12 Hz), beta (12-30 Hz), and gamma (30-50 Hz) frequencies. **(A)** Power spectral density collected from *Syngap1^+/-^* mice was increased compared to *Syngap1^+/+^*mice. **(B)** Delta and theta frequency power were significantly increased in *Syngap1^+/-^* mice. **(C)** Representative power distribution of *Syngap1^+/+^* mice and *Syngap1^+/-^* mice over a 10-minute period illustrates elevated delta and theta power in *Syngap1^+/-^* mice. **(D)** *Syngap1^+/-^* mice exhibited elevated spike train counts. **(E)** *Syngap1^+/-^* mice displayed significantly increased total spike train duration. **(F)** Representative EEG traces of *Syngap1^+/+^* and *Syngap1^+/-^* mice recorded over 10 minutes. Data were analyzed using a two-way ANOVA between genotype and frequency and Sidak’s post hoc analysis where applicable (A,B), or with Student’s t-test (D,E). Data are expressed as mean ± S.E.M. * = *P* < 0.05, ** = *P* < 0.01, *** = *P* < 0.001 **** = *P* < 0.0001.

We characterized four sleep stages and found alterations in *Syngap1^+/-^*mice (**Fig 5A**). We first assessed active wake, during which the subjects are awake and moving measured by EMG signal and activity. *Syngap1^+/-^* mice displayed an increased percentage of time in the active wake stage (**Fig. 5B**; *t*(17) = 3.659, *P* = 0.0019). We then assessed the percent of time in the wake stage where the subjects are awake but not active. *Syngap1^+/-^* mice had a significantly reduced percentage of wake time compared to *Syngap1^+/+^* controls (**Fig. 5C**; *t*(18) = 2.398, *P* = 0.0275). Next, we examined sleep characteristics and found a significant reduction in the percent of slow wave sleep in *Syngap1^+/-^*mice compared to *Syngap1^+/+^* mice (**Fig. 5D**; *t*(17) = 3.060, *P* = 0.0071). Although paradoxical sleep was not significantly affected, *Syngap1^+/-^* mice trended toward a lower percentage of time in paradoxical sleep (**Fig. 5E**; *t*(18) = 1.856, *P* = 0.0800).

**Figure 5.**
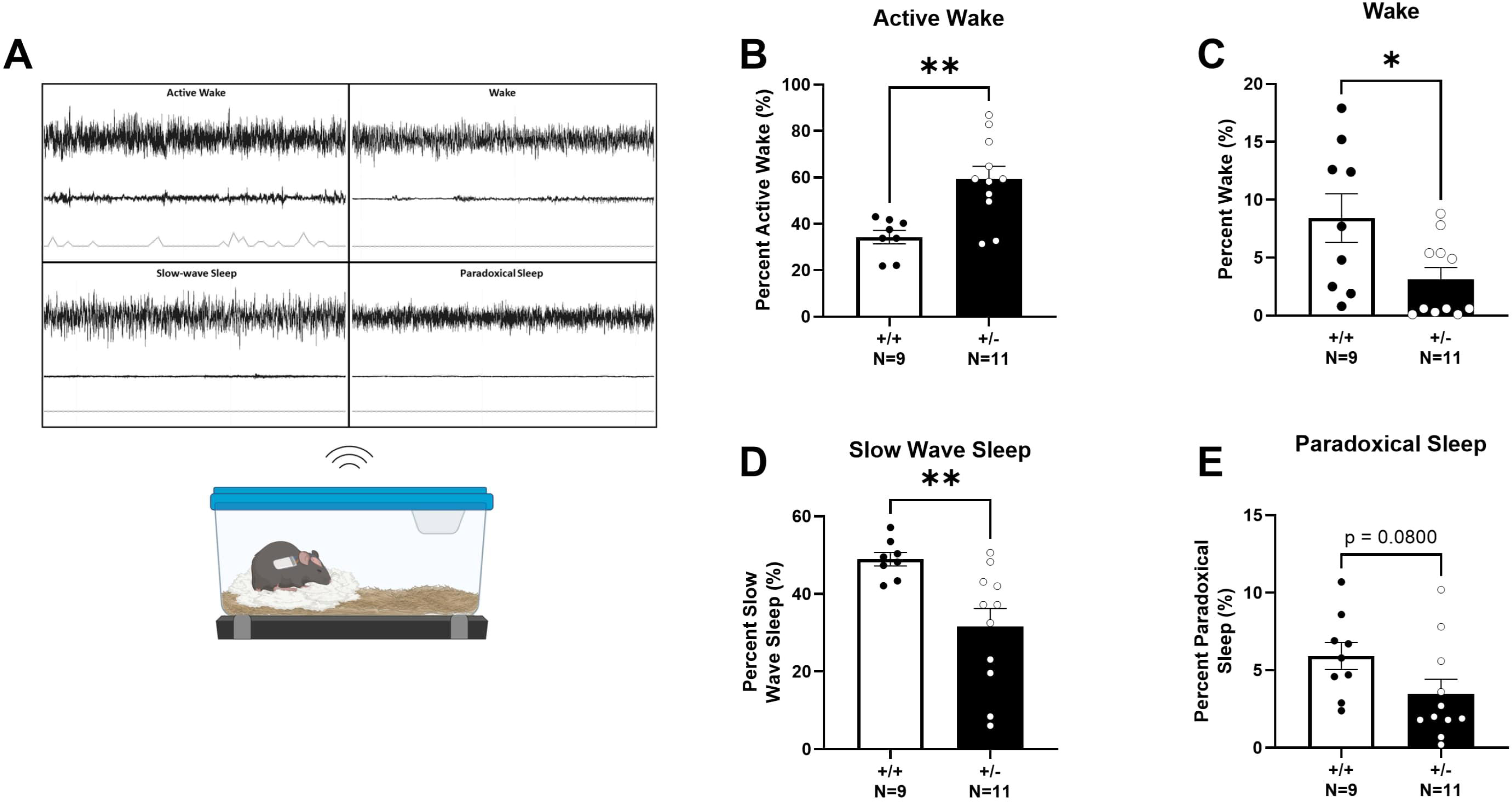
*Syngap1^+/-^*mice displayed altered EEG sleep signatures. **(A)** Schematic of wireless telemetric recording system and representative signals from the four sleep stages. EEG signal is recorded while mice are in their home cage and allowed to move freely. **(B)** Active wake percentage determined from EMG recording and integrated accelerometer when subject displayed movement while awake. *Syngap1^+/-^* mice displayed elevated active wake time. **(C)** Percentage of wake across 72 hours where animals were not actively moving. *Syngap1^+/-^* mice showed reduced wake compared to *Syngap1^+/+^* mice **(D)** Slow wave sleep percentage was decreased in *Syngap1^+/-^* mice. **(E)** Paradoxical sleep percentage from 72 hours of recording trended lower in *Syngap1^+/-^* mice. Data were analyzed using Student’s unpaired t-test. Data are expressed as mean ± S.E.M. * = *P* < 0.05, ** = *P* < 0.01.

### Elevated electrophysiological activity in cultured primary neurons from *Syngap1^+/-^* mice

Electrophysiological activity was assessed in cultured primary neurons from both *Syngap1^+/-^* and *Syngap1^+/+^* mice on high-density multi-electrode arrays. Firing rate from the entire chip area was assessed following an “Activity Scan” of all 26,400 electrodes for 30 seconds (**Fig. 6A**). Electrodes with the highest firing rate were then chosen to perform a “Network Activity Scan” where the activity from 1024 electrodes was recorded simultaneously for five minutes (**Fig. 6B**). Raster plots were generated to visualize bursting events from the simultaneously recorded electrodes. Vertical lines visible on the raster plots indicate coordinated, synchronized bursting activity and were visually elevated in *Syngap1^+/-^* mice (**Fig. 6C**) compared to *Syngap1^+/+^*mice (**Fig. 6D**). Activity was plotted following Gaussian convolution of the spiking data to quantify bursting events for *Syngap1^+/+^*mice (**Fig. 6E**) and *Syngap1^+/-^* mice (**Fig. 6F**). We observed significantly reduced spikes per burst on DIV17 in *Syngap1^+/-^* mice when compared to *Syngap1^+/+^* mice via Sidak’s post hoc analysis following two-way ANOVA (**Fig. 7A**; *F*(1, 13) = 11.56, *P* = 0.0418) and DIV21 (**Fig. 7A**; *P* < 0.0001). Inter-burst interval was significantly reduced in the Syngap1^+/-^ mice on DIV21, DIV27, and DIV29 when analyzed with Sidak’s multiple comparison test following two-way ANOVA (**Fig. 7B**; *F*(1, 76) = 20.33, *P* = 0.0068 (DIV21), *P* = 0.0010 (DIV27), *P* = 0.0035 (DIV29)). The reduced inter-burst interval in the *Syngap1^+/-^*mice aligns with the increased number of bursts that we observed on DIV21, DIV27, DIV29, and DIV35 when compared to neurons from *Syngap1^+/+^* mice following post hoc analysis (**Fig. 7C**; *F*(1, 78) = 32.86, *P* = 0.0129 (DIV21), *P* = 0.0003 (DIV27), *P* = 0.0004 (DIV29), *P* = 0.0013 (DIV35)). We also measured an increased burst duration on DIV29 in *Syngap1^+/-^*neurons compared to *Syngap1^+/+^* neurons (**Fig. 7D**; *F*(1, 64) = 4.166, *P* = 0.0229). These data indicate that neurons from *Syngap1^+/-^* mice exhibit burst firing more frequently than neurons from *Syngap1^+/+^* mice at comparable time points.

**Figure 6.**
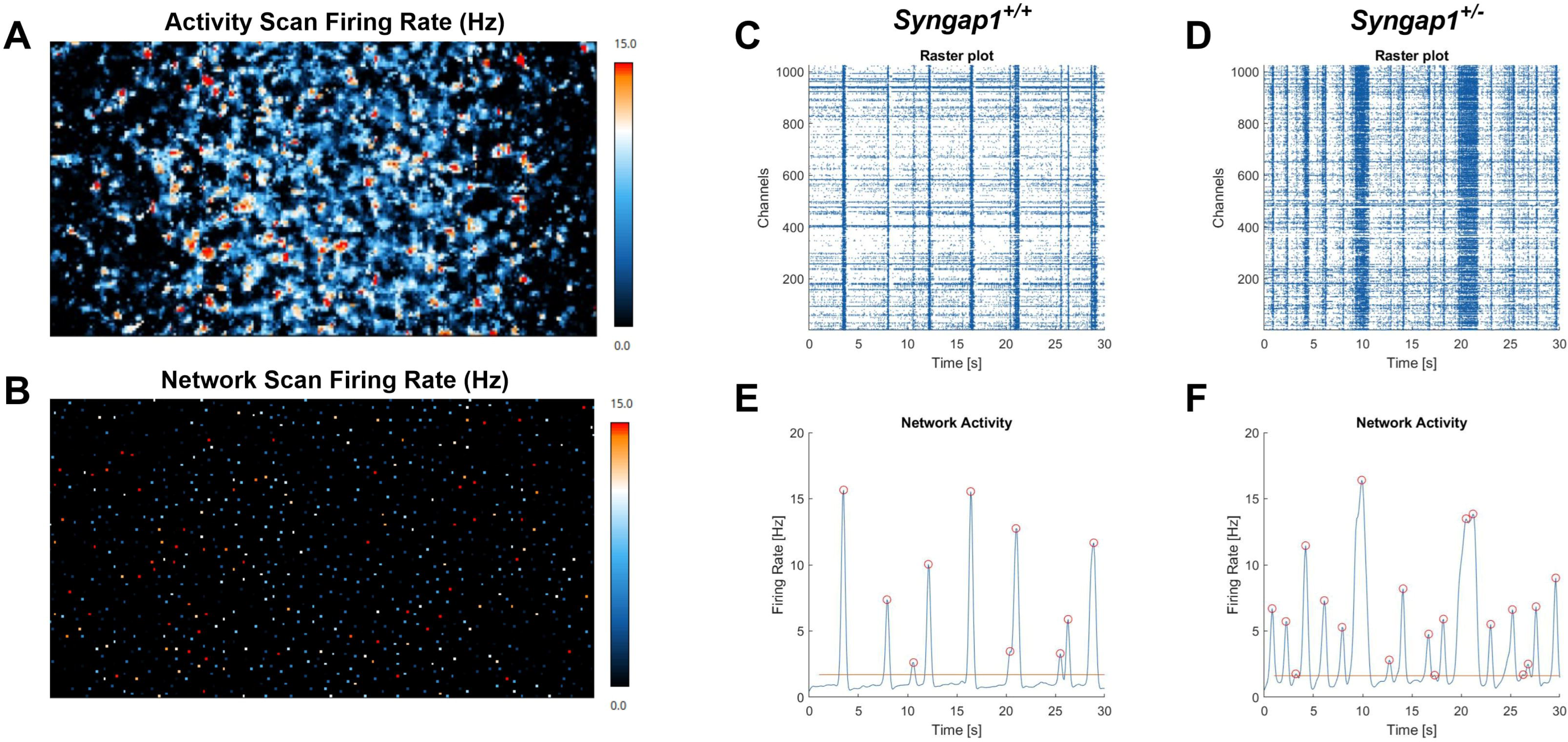
Primary neurons from *Syngap1^+/-^*mice displayed increased network firing activity when measured with high-density multi-electrode arrays (HD-MEAs). **(A)** Firing rate from the entire chip area. **(B)** 1024 electrodes with the highest firing rates were chosen to conduct a “Network Activity Scan”. **(C)** Representative raster plot and **(E)** network activity plot of *Syngap1^+/+^* primary neurons. **(D)** *Syngap1^+/-^* raster plot and **(F)** network activity plot exhibit increased activity. 1024 electrodes were recorded simultaneously and plotted on the y-axis of raster plots.

**Figure 7.**
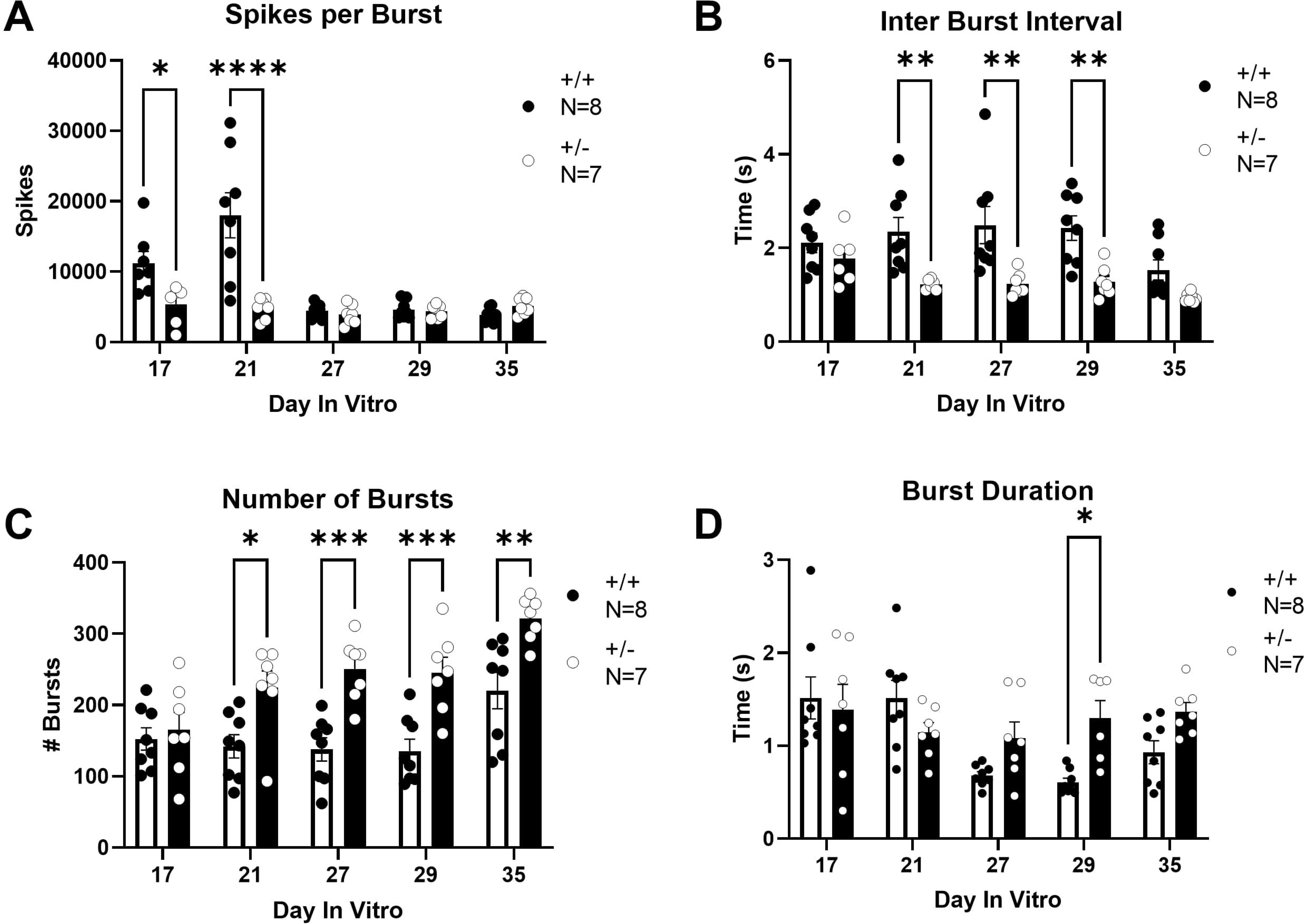
*Syngap1^+/-^* mice displayed increased bursting activity when measured on HD-MEAs. **(A)** Spikes per burst in *Syngap1^+/+^*and *Syngap1^+/-^* mice. **(B)** Inter burst interval (IBI) was measured as time between burst peaks. A reduced IBI was observed in *Syngap1^+/-^* animals on DIVs 21, 27, and 29. **(C)** The total number of bursts over the five-minute recording was compared. *Syngap1^+/-^* mice exhibited an increased number of bursts on all days after DIV21. **(D)** *Syngap1^+/-^*mice had a longer burst duration on DIV29 when compared to *Syngap1^+/+^*mice. Data were analyzed using a two-way ANOVA. Data are expressed as mean ± S.E.M. * = *P* < 0.05, ** = *P* < 0.01, *** = *P* < 0.001 **** = *P* < 0.0001.

## Discussion

A myriad of NDDs result from missing/mutated proteins of the postsynaptic density (PSD), including Shanks,^26, 31^ Homers,^32–34^ mGluRs,^35^ and SynGAP1. SynGAP1 is the major neuronal specific RasGAP that binds to the PSD-95, a major scaffolding protein of the PSD. SynGAP1 is localized to excitatory synapses and is one of the most highly abundant components of the PSD.^1, 2^ The PSD is composed of densely organized proteins adjacent to the post-synaptic membrane comprised of anchoring cell surface proteins, scaffold proteins, neurotransmitter receptors, and cell-adhesion molecules. SynGAP1 is primarily expressed in the brain, thus it is not surprising that perturbations in SynGAP1 expression result in a pathological NDD.^2, 3, 36^

We observed rigorous, robust behavioral alterations by hyperactivity in the open field arena, impairments in novel object recognition, and reduced anxiety-like behavior in *Syngap1^+/-^* mice compared to *Syngap1^+/+^* littermate controls, using a unique F1 generation hybrid mouse as the background strain. Advantages of this F1 include the hybrid vigor as a background for deleterious mutations and elimination of the resistance to typical seizure induction methods, a documented phenotype of the classic congenic B6J background strain. ^23, 37, 38^ A comprehensive behavioral battery performed at the neurobehavioral phenotyping laboratory at the Jackson Laboratory found the 129S1-C57BL/6J F1 hybrid to be behaviorally indistinguishable from C57BL/6J. Identical observations regarding lethality when a synaptopathy is on a congenic C57BL/6J background yet excellent thrive ability on 129S1 were observed in *Shank1* mutant mice.^31, 39, 40^ Previous research using the F1 generation of hybrid mice allowed for comprehensive behavioral analysis of *Shank1* model mice, as well as the *Syngap1* mutant mice described herein. Hybrid mice often allow for the detection and expression of polygenic diseases by presenting a broader array of responses to various stresses, thus providing an approximate control for some genetically engineered strains. In our case, investigating the construct valid *Syngap1* mutant mice on C57BL/6J or 129S1 background strains alone, would have prevented data with ample power for statistical conclusions and/or reduced the clarity of our conclusions.

Currently, only a single other study investigated the epileptogenic, and sleep biomarkers of SYNGAP1R-ID. In a 3-year-old child with SYNGAP1R-ID and the *Syngap1* mutant mice on a C57BL/6J background, which is notorious for seizure resistance,^23, 41^ progressive changes in the sleep architecture over 24 hours were reported. Over 24-hours of nocturnal rhythms, WT mice at PND60 and PND120 had less awake time than *Syngap1^+/-^*mice at similar ages.^42^ Our report extends this result and debuts the generalization of sleep alterations to a broader background strain, as well as the use of 4 sleep stages via EEG/EMG over the wake and NREM stages used in the Sullivan work, in a preclinical mouse model of SYNGAP1R-ID. Sleep disturbances are a significant translational phenotype in synaptopathies, such as Phelan McDermid Syndrome (Shank3) and SYNGAP1-ID.^43^ A one-of-a-kind SynGAP1 rat model exists, generated via a collaborative effort between the Simon’s Foundation and the University of Edinburgh. These rats were generated on a Long Evans background and published under the *Syngap^+/^*^Δ*−GAP*^ nomenclature. In the rat study, sleep was analyzed via video-EEG and an automated program for sleep spindles.^44^ Sleep abnormalities were mostly uncorrelated to the electrophysiological signature of absence seizures, spikes, and spike wave discharges. Visual brain sleep state scoring blind to animal genotype was performed by assigning 5 s epochs to non-rapid eye movement sleep (NREM), rapid eye movement sleep (REM) or wake. Scoring criteria for visual classification was based on accelerometer and EEG characteristics like the methodology herein, however we also utilized an EMG signal. Using group sizes of 4 per genotype (without the mention of sex), the electro graphical correlate of absence seizures, spike, and wave discharges (SWDs), were identified visually and analysis was confirmed with an automated algorithm. Briefly, SWDs are characterized by periodic high-amplitude oscillations in the theta band between 5 and 10 Hz which correlates with a spontaneous stop in animal movement. Power spectral analysis was performed that identified harmonic peaks. Via this described methodology of sleep stages, it was reported that Syngap^+/Δ−GAP^ rats spent an equivalent percentage of time in all states when compared with wild-type littermate controls. Tailored further examination showed that wake and NREM bouts, with REM bouts remaining unchanged coinciding with increased average bout duration during wake and NREM, as well as no difference in REM bout duration. Therefore, only during 6 h recordings, *Syngap*^+/Δ−GAP^ rats display an abnormal sleep state distribution. All data were classified by individual 5 s recording epochs from those previous recordings as NREM sleep, REM sleep or wake. While direct comparison of our data to this earlier work is complicated given the vastly different methodologies, species, and EEG acquisition time, the consensus of these 3 studies is that sleep will be a powerful translational predictor in a future clinical trial for SYNGAP1R-ID^44^ as has been demonstrated in other rare genetic NDDs.^45–47, 48, 49^

EEG recordings in neurodevelopmental disorders show potential to identify clinically translatable biomarkers to both diagnose and track the progress of novel therapeutic strategies, as well as providing insight into underlying pathological mechanisms. When spontaneous recurring seizures were observed by visual scoring of a 24 h video EEG, few seizures were observed in the *Syngap1^+/-^* mice until PND120 (4 months of age) a time at which EEG seizures greatly increased.^42^ Our data corroborate work using the indices of spiking and spike trains, analyzed using the same methods as Baylor Neurology.^50, 51^ We also utilized the oscillatory power to obtain power spectral densities of each frequency wave, discovering greater overall power in *Syngap1^+/-^*mice, and elevated delta and theta power. Elevated delta power is being used as a biomarker in clinical trials for other NDDs, such as Angelman Syndrome clinical trials.^25, 51–55^ Increased spike trains during *in vivo* EEG show similar patterns of activity to the increased numbers of bursts and shorter latency bursts using micro-electrode arrays (MEA) on *Syngap1^+/-^*primary neurons. This report debuts this innovative technology in this genetic mouse model as we have discovered a functional physiology outcome that bridges our *in vitro* studies to our *in vivo* results. Moreover, this technology can be used with other cell types and record from neural stem cells generated from human iPSCs which could serve as another translational bridging study.^28^ Using human cells, high density MEAs, such as those described herein, have produced rigorous, reliable, reproducible findings of genetically generated disease states and by observing expected firing changes with pharmacological tools.^27^ Continuous and long-term recordings of the circuit dynamics was not possible due to phototoxicity and the large number of challenges in maintaining seals for classical patch clamping techniques.^29^ Here, we utilize a high-density microelectrode array containing 26,400 electrodes and can simultaneously record 1024 discrete electrodes for label-free, comprehensive, and detailed electrophysiological live neuronal cell recording of over 3-4 weeks in culture.^27, 28^

As a negative regulator of excitatory neurotransmission, overexpression of *Syngap1* results in a dramatic loss of synaptic efficacy as well as enhanced synaptic transmission following SynGAP1 disruption by RNA interference.^4^ This work is vital, proving that SynGAP1 levels are modifiable and SynGAP1-deficient synapses are not immutable. Added hope comes from recent work that illustrates that Syngap1 is a downstream target of MAPK interacting protein kinases 1 and 2 (Mnk1/2),^56^ which regulate a plethora of functions, presumably via phosphorylation of substrates, including eukaryotic translation initiation factor 4E (eIF4E). Reducing Syngap1 levels reversed behavioral learning and memory deficits in a Mnk ½ double knockout mouse model, leading to the novel suggestion that the Mnks–Syngap1 axis regulates memory formation and functional outcomes.^56^

Key questions that remain for all NDDs include age of restoration, and “critical windows”. Many have described that rescues are possible as adults in Fragile X,^57, 58^ Rett,^59, 60^ Phelan-McDermid,^61^ and Angelman Syndromes.^18, 20, 62^ However, others have argued that only intervention in early life reverses behavioral phenotypes in some NDDs.^63^ In fact, prenatal intervention theories are being explored.^64^ Earlier work with SynGAP1 illustrated hardwiring of neural circuitry that manifest as lifelong impairments.^65, 66^ Nonetheless, crucial to this work, re-expression of *Syngap1* by genetic reversal exhibits a complete alleviation of electrophysiological and cognitive behavioral phenotypes in a genetic inducible mouse model,^67^ suggesting nearly full expression of a second allele of *Syngap1* is required for alleviation of regulatory issues, regarding potency, target engagement, and PK/PD for transability. Our laboratory is currently assessing nuanced SynGAP1 alterations using an ELISA assay over semi-quantitated Western blots or RNA levels only, which can lack predictability. Genetic reversal was illustrated only when re-expression was localized to glutamatergic neurons, which contribute significantly but not in isolation to the phenotypes reported herein. It is currently not known if other neuronal subtypes are also sufficient to drive the reported abnormalities in these mice, however our current targeted therapeutics under investigation in this construct valid model are addressing that exact question.

## Supplementary Material

Supplementary material is available at *Brain* online.

## Supplementary Results and Supplemental Figure Legend

### Hyperactivity and reduced anxiety like phenotype

Anxiety -like activity was assessed using the elevated plus maze and the light-dark box tasks. An increased percentage of time spent in the open arms of the elevated plus maze was observed in the Syngap1+/-mice when compared to Syngap1+/+ mice using a student’s unpaired t-test (**Fig. S1A**; *t*(46) = 2.471, *P* = 0.0172) indicating reduced anxiety-like behavior. Syngap1+/-mice also showed heightened open arm entries (**Fig. S1B**; *t*(46) = 4.104, *P* = 0.0002) as well as total transitions between open and closed arms in the elevated plus maze when analyzed with an unpaired t-test (**Fig. S1C**; *t*(46) = 5.240, *P* < 0.0001). Similar elevated total entries were observed in the light-dark task with the Syngap1+/-mice showing a significantly increased number of total entries into the light space when compared to Syngap1+/+ mice (**Fig. S1D**; *t*(39) = 4.481, *P* < 0.0001).

**Figure S1.**
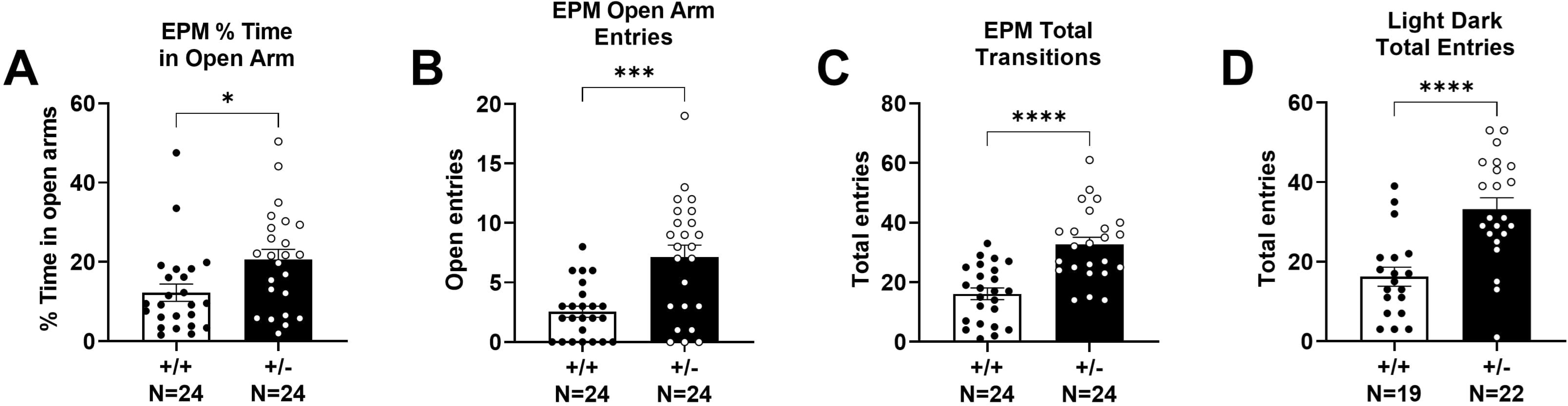
Reduced anxiety-like behavior and hyperactivity when assessed in the elevated plus maze (EPM) and the light-dark conflict task. **(A)** In the EPM, percent time spent on the open arms, **(B)** the total number of open arm entries, and the **(C)** total number of transitions were elevated in *Syngap1^+/-^*mice when compared to *Syngap1^+/+^* mice. **(D)** In the light-dark task, total transitions between chambers was elevated in the *Syngap1*^+/-^ mice. Data are expressed as mean ± S.E.M. * = *P* < 0.05, *** = *P* < 0.001, **** = *P* < 0.0001 when analyzed with a student’s unpaired t-test.

**Figure S2.**
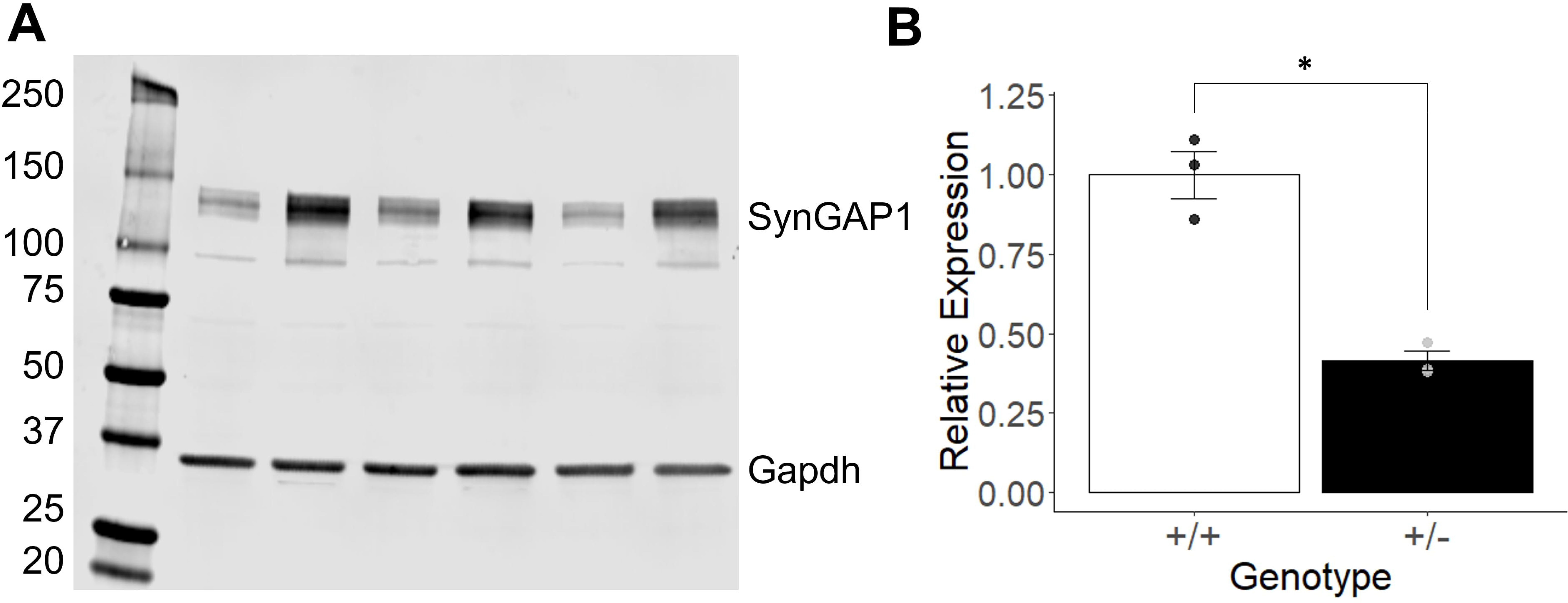
Unedited western blot of SynGAP1 and Gapdh protein expression in *Syngap1^+/+^* (WT) and *Syngap1^+/-^* (HT) mice at PND42. **(A)** A significant decrease in expression of SynGAP1 was observed in the *Syngap^+/-^* mice. Bands not at 140 kDa or 37 kDa are non-specific and do not show significant changes between genotypes. **(B)** Quantification of SynGAP1 protein expression using Gapdh expression as normalization. SynGAP protein expression was decreased to 41% of WT expression. Data was analyzed using a Student’s t-test and is expressed as mean ± S.E.M. \**P* = 0.0051. (WT *N* = 3, HT *N* = 3)

